# Small molecule STAT3/5 inhibitors exhibit therapeutic potential in acute myeloid leukemia and extra-nodal natural killer/T cell lymphoma

**DOI:** 10.1101/2023.10.12.561697

**Authors:** Daniel Pölöske, Helena Sorger, Anna Schönbichler, Elvin D. de Araujo, Heidi A. Neubauer, Anna Orlova, Sanna H. Timonen, Diaaeldin I. Abdallah, Aleksandr Ianevski, Heikki Kuusanmäki, Marta Surbek, Christina Wagner, Tobias Suske, Martin L. Metzelder, Michael Bergmann, Maik Dahlhoff, Florian Grebien, Roman Fleck, Christine Pirker, Walter Berger, Emir Hadzijusufovic, Wolfgang R. Sperr, Lukas Kenner, Peter Valent, Tero Aittokallio, Marco Herling, Satu Mustjoki, Patrick T. Gunning, Richard Moriggl

## Abstract

The oncogenic transcription factors STAT3, STAT5A and STAT5B are essential to steer hematopoiesis and immunity, but their enhanced expression and activation drives the development or progression of blood cancers. Current therapeutic strategies focus on blocking upstream tyrosine kinases, but frequently occurring resistance often leads to disease relapse, emphasizing the need for more targeted therapies. Here we evaluate JPX-0700 and JPX-0750, which are STAT3/5-specific covalent cysteine binders that lead to growth arrest of acute myeloid leukemia (AML) and natural killer/T cell lymphoma (NKCL) cell lines *in vitro* and *in vivo*, as well as reduce cell viability of primary AML blasts *ex vivo*. Our non-PROTAC small molecular weight degraders selectively reduce STAT3/5 activation and total protein levels, as well as downstream target oncogene expression, exhibiting nanomolar to low micromolar efficacy. We found that both AML and NKCL cells hijack STAT3/5 signaling through either upstream activating mutations in tyrosine kinases, activating gain-of-function mutations in STAT3, mutational loss of negative STAT regulators, or genetic gains in anti-apoptotic, pro-proliferative or epigenetic-modifying STAT3/5 targets. Moreover, we have shown synergistic inhibitory action of JPX-0700 and JPX-0750 upon combinatorial use with approved chemotherapeutics (doxorubicin, daunorubicin, cytarabine), epigenetic enzyme blocker vorinostat, tyrosine kinase inhibitor cabozantinib or BCL-2 inhibitor venetoclax. Importantly, JPX-0700 or JPX-0750 treatment reduced leukemic cell growth in human AML/NKCL xenograft mouse models without adverse side effects. These potent small molecule degraders of STAT3/5 could propel further clinical development for use in AML and NKCL patients.

## Introduction

STAT3 and the closely related STAT5A and STAT5B transcription factors are encoded by three different genes on human chromosome *17q*. Importantly, *17q* gain is genetically the most frequent chromosomal abnormality in cancer, observed in 2.5% of all cancer cases (1), which can promote overexpression of STAT3/5 as shown in cutaneous T cell lymphoma (2) and prostate cancer (3). All three STAT3/5 proteins play key effector roles upon JAK kinase signaling downstream of a plethora of cytokines, or they can be directly activated upon growth factor receptor engagement (4). STAT3/5 proteins regulate vital cellular processes in hematopoietic cells, such as proliferation, survival, differentiation, metabolism or chromatin remodeling (5, 6). Enhanced STAT3/5 expression, gain-of-function (GOF) mutations or prolonged activation frequently occurs in cancer and are associated with higher sensitivity to microenvironment stimuli and cancer proliferation or survival (7, 8). Interestingly, among the many different blood cancers, two classes have a particularly strong *STAT3/5* gene expression signature, namely acute myeloid leukemia (AML) and extra-nodal NK/T cell lymphoma (NKCL) (9, 10). AML is associated with increased STAT3/5 signaling due to upstream driver mutations in tyrosine kinases (TK) such as *FLT3,* or oncogenic GTPase hyperactivation such as NRAS or RHOA, promoting either nuclear shuttling of STAT5 or STAT3/5-mediated alterations to cellular metabolism (11, 12). FLT3-internal tandem duplication (ITD) rearrangements lead to enhanced STAT5 activation, due to additional docking sites for STAT5 on the receptor (from 8 to >800 bp, with longer rearrangements resulting in more STAT5 docking sites and poorer cancer prognosis), boosting STAT5-induced gene transcription (13). Furthermore, GOF variants of STAT3/5 are common in lymphoid malignancies, such as NKCL, in which patients frequently harbor GOF mutations in *STAT3* (∼30%), *STAT5A/B* (∼10%) and *JAK3* (∼10%) in addition to mutations in tumor suppressors *PRDM1* (∼40%), *TP53* (∼10%), chromatin modifiers *KMT2D* (∼15%), *KMT2C* (∼15%) and the family of DEAD-box helicases (∼45%) (14–16). Both AML and NKCL diseases are difficult to control with conventional therapy.

Past efforts to inhibit STAT3/5 signaling focused on targeting upstream TK, therefore indirectly blocking STAT3/5 (17, 18). However, tumor cells frequently develop resistance to conventional cytostatic agents. Mutations within the kinase driver itself, enhanced expression of STAT3/5 or enhanced expression/mutations in downstream effector molecules, such as c-MYC, n-MYC, D-type Cyclins, AXL or BCL-2 family members (19, 20), often lead to therapy failure in AML or NKCL patients (21, 22). Therefore, despite different mechanisms of STAT3/5 hyperactivation and dependence, the dual targeting of oncogenic STAT3/5 could be a novel and promising therapeutic strategy in myeloid and lymphoid malignancies.

Using AML and NKCL as model systems, we investigated the potential of directly targeting STAT3/5 as a novel and more effective therapeutic strategy to combat cancer growth. We employ two covalent small molecule STAT3/5 pathway inhibitors, JPX-0700 and JPX-0750, which we have shown to directly bind and destabilize both STAT3 and STAT5 proteins *in vitro*, and eliminate both phosphorylated and total STAT3/5 in cells (23, 24). Here, we screened a large panel of blood cancer cell lines and demonstrated sensitivity of particularly NKCL and AML cells to these compounds, both of which display high levels of STAT3/5 signaling. Profiling genetic hallmarks, we reveal chromosomal gains in pro-proliferative/anti-apoptotic STAT3/5 targets in NKCL, including *MYC, MYCN, BCL2L1, MCL1, DNMT3A* and *DNMT3B,* or lost negative regulators such as *TP53, CDKN2A* and *PTPN2,* indicating further amplification of STAT3/5 oncogenic signaling. We show that these inhibitors facilitate rapid STAT5 degradation, with a relatively slower STAT3 degradation profile, leading to the downregulation of oncogenic STAT3/5 targets *in cellulo*. AML and NKCL cancer cells and primary AML blasts showed significant loss of viability in the presence of the STAT3/5 inhibitors, and tumor growth was reduced in subcutaneous xenograft models in the absence of toxic side effects. Moreover, we found synergistic cytotoxic effects when combining these STAT3/5 inhibitors with other clinically approved drugs used in AML or NKCL treatment.

## Results

### JPX-0700 and JPX-0750 exhibit marked potency in AML and NKCL cell lines

Due to IL-2 dependence and frequent *STAT3/STAT5A/STAT5B* mutations in NKCL, or general tyrosine kinase inhibitor (TKI) resistance mechanisms in AML, we hypothesized that the simultaneous inhibition of STAT3/5 might decrease the oncogenic signaling cascade induced by STAT3/5 activation and block proliferation and survival (**Supplementary Figure S1A, B, Supplementary Table S1**). We aimed to evaluate the efficacy of JPX-0700 and JPX-0750 on several human blood cancer cell lines and correlate their sensitivity to the activation-or overall STAT3/5 protein levels. For this, we analyzed FLT3-ITD^+^ and FLT3-ITD^-^ AML cell lines, NKCL cell lines, representing NK large granular lymphocytic leukemia (NK-LGL), extra-nodal NK/T cell lymphoma, nasal type (ENKL), aggressive NK cell leukemia (ANKL) and NK large lymphocytic leukemia, and T cell lymphoma (TCL) cell lines, namely anaplastic large cell lymphoma (ALCL) and cutaneous T cell lymphoma (CTCL) cell lines. Control cells used in this study represent a lung adenocarcinoma (A549) and human fibroblast cell line (NHDF). Cell viability assessment upon 72 h of JPX-0700 and JPX-0750 inhibitor treatment showed a 10-fold increase in potency in FLT3-ITD-driven AML cell lines, MV4-11 [0.17 µM and 0.10 µM] and MOLM-13 [0.35 µM and 0.19 µM], in comparison to the previously published STAT5 inhibitor AC-4-130 (10) (**Figure 1A**). NKCL and TCL cell lines revealed low nM to µM IC_50_ values across the cell line panel for JPX-0700 [0.29 – 2.59 µM] and JPX-0750 [0.47 – 2.83 µM]. Notably, NKCL were the most sensitive lymphoid cancer in our analysis, responding slightly better towards JPX-0700 treatment. The investigated blood cancer cell lines displayed either predominantly activated STAT5 (MV4-11, MOLM-13, NKL), STAT3 (SU-DHL-1), or activation of STAT3/5 (SNK-6, NK-YS, KHYG-1, Mac-2A), highlighting the importance of STAT3/5 signaling in these malignancies (**Figure 1B**). Interestingly, the IC_50_ values increased proportionally with higher STAT3/5 protein levels in NKCL/TCL cell lines, while lower IC_50_ values in AML significantly correlated with lower total protein and STAT3/5 activation levels (**Figure 1B, Supplementary Figure S1C, B**). A549 and NHDF cells showed no significant STAT3/5 phosphorylation and were, due to their high IC_50_ values, regarded as predominantly resistant towards STAT3/5 inhibition.

**Figure 1.**
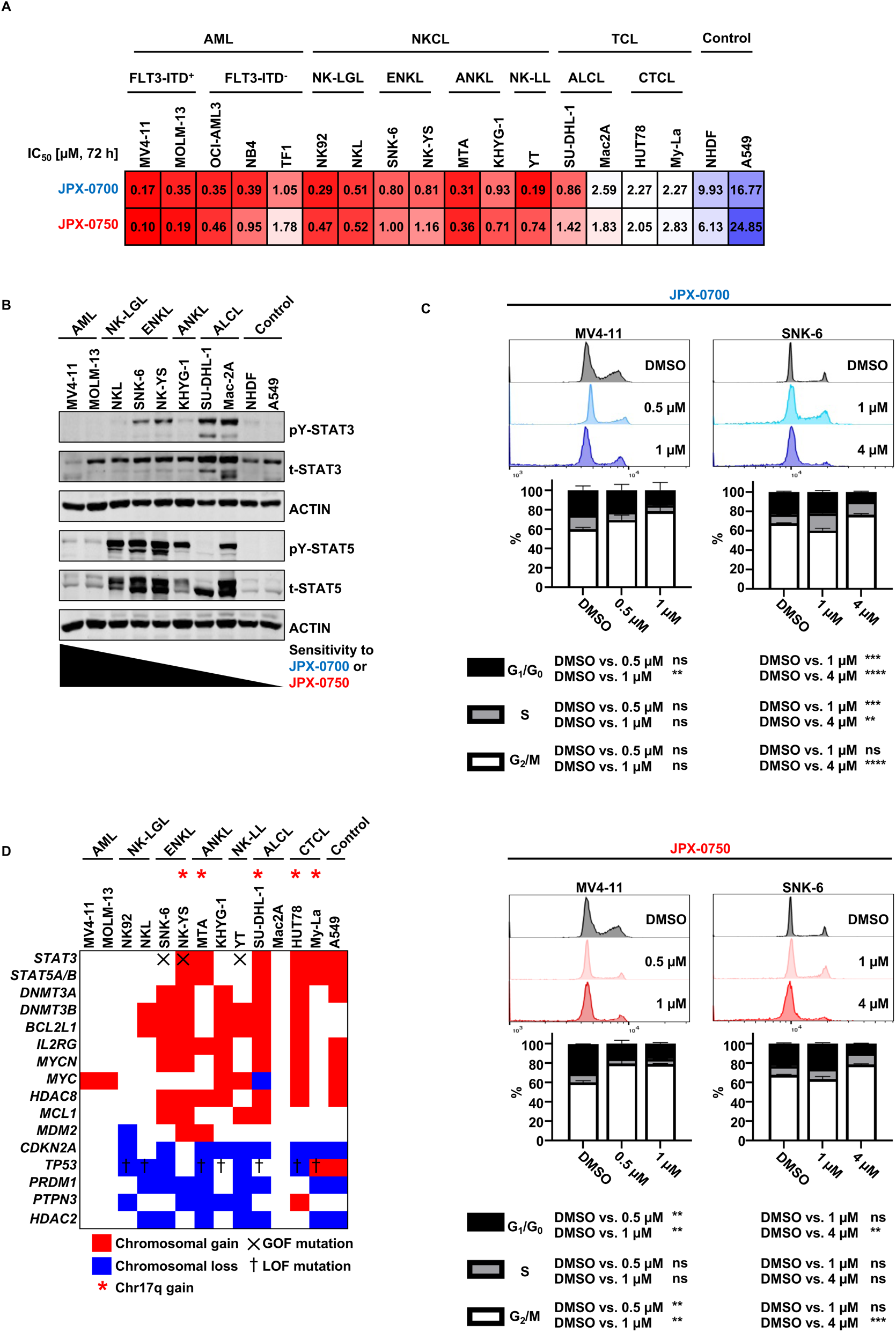
Treatment of AML and NKCL cell lines with selective STAT3/5 inhibitors JPX-0700 and JPX-0750 leads to G1/G0 cell cycle arrest, inhibition of cell proliferation and cell death. A, Heatmap of IC50 values calculated from drug response analysis using CellTiter-Blue viability assays upon 72 h treatment with dual STAT3/5 inhibitors, JPX-0700 or JPX-0750. One representative, out of three independent experiments performed in triplicates, is shown (N=3). B, Protein extracts from cell lines derived from either AML or various subtypes of NKCL and TCL malignancies were immunoblotted for total and phospho-Tyr (705)-STAT3 and total and phospho-Tyr (694/699)-STAT5A/B. ACTIN served as loading control (N=2). C, Cell cycle analysis of MV4-11-AML and SNK-6-ENKL cells after 72 h treatment with JPX-0700 or JPX-0750. One representative, out of two independent experiments is shown (N=2). D, aCGH analysis showing most frequent chromosomal mutations of genes involved in STAT3/5 signaling in NKCL, TCL, AML cell lines. Chromosomal gains are marked in red and losses in blue, X: gain-of-function mutation, †: loss-of-function mutation, *: Chr17q gain. Data represent the mean ± SEM. *p < 0.05, **p < 0.01, and ***p < 0.001, by 2-way ANOVA with Bonferroni’s correction. Abbreviations: aCGH=array comparative genome hybridization, ALCL=anaplastic large cell lymphoma, AML=acute myeloid leukemia, ANKL=aggressive NK cell leukemia, CTCL=cutaneous T cell lymphoma, ENKL=extra-nodal NK/T cell lymphoma, nasal type, NKCL=NK cell lymphoma/leukemia, NK-LGL=NK large granular lymphocytic leukemia, NK-LL=NK large lymphocytic leukemia, TCL=T cell lymphoma/leukemia

We observed an arrest of AML and NKCL cells primarily in the G_1_ phase and a depletion of cells in the G_2_/M phase upon JPX-0700/0750 treatment, indicating a growth arrest. Furthermore, drug-induced cell cycle arrest was accompanied by poly ADP-ribose polymerase (PARP) and Caspase-3 cleavage, key mediators of DNA damage/apoptosis pathways (**Figure 1C, Supplementary Figure S1D**). To evaluate the *in cellulo* target specificity of these compounds, we used the murine IL-3 dependent cell line Ba/F3, modified to overexpress either STAT3, STAT5A or STAT5B. Treatment of cells with JPX-0700 or JPX-0750 for 16 h led to loss of STAT3/5 activation and degradation of STAT3/5 total protein (**Supplementary Figure S1E-J**). As expected, higher concentrations of inhibitors were required to degrade the STAT3/5 protein in the overexpressing cell lines compared with endogenous protein in the parental Ba/F3 cells.

To explain the heightened susceptibility towards STAT3/5 inhibition of AML and NKCL cell lines, we performed a genomic study using array comparative genome hybridization (aCGH) analysis to link cell line sensitivity to genomic aberrations impacting the STAT3/5 signaling pathway. We extended previous findings of the genetic profiling of NKCL (9, 25), compared them to AML profiles (**Supplementary Table S1**), and observed recurrent copy number (CN) gains in *STAT3, STAT5A/B*, *BCL2L1*, *MCL1*, *IL2RG*, *MYCN*, *MYC*, *HDAC8*, *MDM2*, *DNMT3A* and *DNMT3B*. Moreover, frequent chromosomal losses in tumor suppressors, namely *PTPN3*, *TP53*, *CDKN2A* and *PRDM1,* were also evident (**Figure 1D**). Interestingly, recurrent *HDAC2*-loss was apparent, which could indicate a potential tumor suppressive function, warranting further study in NKCL.

### Small molecule inhibitors induce programmed cell death and block STAT3/5 downstream target gene expression in AML and NKCL

Analyzing the mode of action of cell killing revealed that the small molecule inhibitors reduced cell viability by inducing cell death, as demonstrated by Annexin V/Propidium iodide staining in the FLT3-ITD driven MV4-11 and MOLM-13 AML cells and the NKCL cell lines SNK-6 and NK-YS, upon 24 h treatment with JPX-0700/0750 (**Figure 2A, Supplementary Figure S2A**). Measuring the kinetics of cell death induction revealed that both JPX-0700 and JPX-0750 are more efficacious in MV4-11 cells [57% and 36% live cells], compared to SNK-6 cells [69% and 69% live cells] at 1 µM concentration, which is most likely attributable to lower STAT3/5 levels and higher sensitivity towards STAT3/5 inhibition (**Figure 1B, Supplementary Figure S1C**).

**Figure 2.**
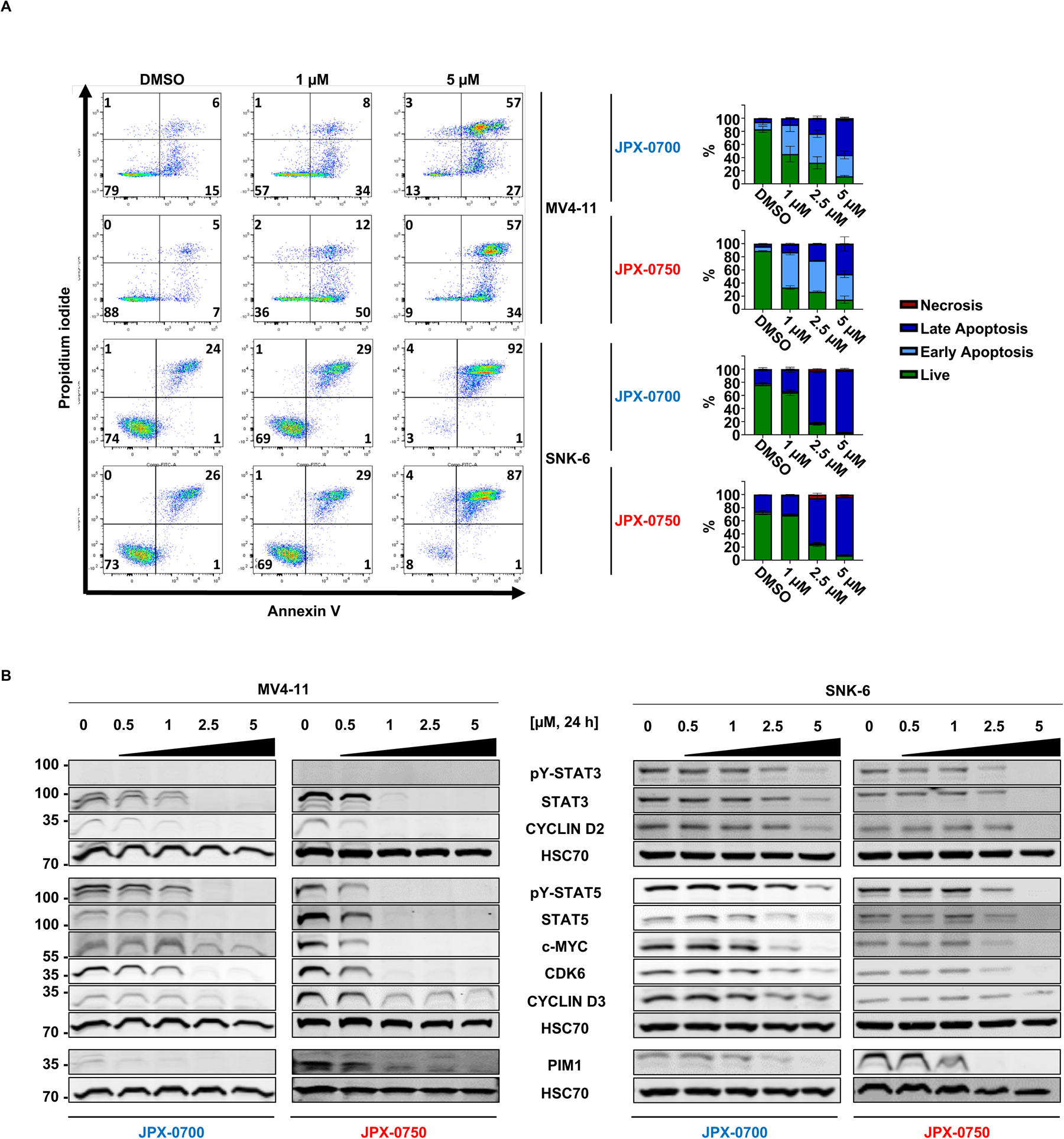
Small molecule STAT3/5 inhibitors reduce downstream STAT3/5 targets such as important oncoproteins thereby inducing cell death and arresting AML and NKCL cell proliferation. A, MV4-11 and SNK-6 cells were treated with different concentrations of JPX-0700, JPX-0750 or DMSO in a dose-dependent manner for 24Lh. Apoptotic cells were detected by Annexin-V/Propidium iodide staining (N=2, representative dot plots and quantification bar graphs are shown). B, Inhibitor treatment was carried out with increased dose escalation for 24 h in MV4-11 and SNK-6 cell lines. Cells were immunoblotted for total and phospho-Tyr (705)-STAT3 and total and phospho-Tyr (694/699)-STAT5A/B. STAT3/5 target gene products c-MYC, PIM1, as well as proteins regulating the cell cycle, such as CYCLIN D2 and D3, and CDK6, were probed by Western blotting. HSC70 served as loading control (N=2, representative experiments are shown).

Next, we performed immunoblotting of STAT3/5, and their target gene products, to confirm a downstream impact on the signaling pathway in AML and NKCL cell lines. We focused on target genes that encode oncoproteins involved in cell proliferation and survival, such as c-MYC, PIM1 and proteins regulating the cell cycle, such as CYCLIN D2 and D3, and CDK6 (2, 10, 26). JPX-0700 and JPX-0750 degrade STAT3/5 proteins effectively in the AML and NKCL cells (**Figure 2B**). Degradation was stronger in AML cells at 1 µM after 24 h treatment, in line with the more efficient induction of cell death in these cells. Importantly, JPX-0750 completely abrogated c-MYC, CYCLIN D2 and CDK6 levels at 1 µM, but surprisingly CYCLIN D3 and PIM1 were only slightly downregulated, suggesting either a longer protein half-life or alternative routes of activation of these oncogenes. CYCLIN D3 and PIM1 downregulation was stronger upon 2.5 µM JPX-0700 treatment in the NKCL cell line after 24 h treatment (**Figure 2B, Supplementary Figure S2B**). In summary, JPX-0700/0750 are potent, cell death-inducing compounds that inhibit STAT3/5 and abrogate downstream oncoprotein nodes in AML and NKCL cells

### JPX-0700 and JPX-0750 are well tolerated by mice and effectively suppress AML and NKCL xenograft growth in vivo

To assess whether JPX-0700 and JPX-0750 can inhibit AML and NKCL tumor growth *in vivo*, we established xenograft models using the MV4-11 and the SNK-6 cell lines. Leukemic cells were subcutaneously injected into both flanks of NOD.Cg-Prkdcscid Il2rgtm1Wjl Tg(CMV-IL3,CSF2,KITLG)1Eav/MloySzJ NOD scid gamma (NSGS) mice. Based on our *in vitro* IC_50_ data (**Figure 1C**), we chose to treat SNK-6 xenografts with JPX-0700 and MV4-11 xenografts with JPX-0750. JPX-0700 or JPX-0750 (5 mg/kg) was intraperitoneally administered daily for 19 or 12 days, respectively, beginning once tumors were palpable at 29 (SNK-6) or 8 days (MV4-11) post-injection (**Figure 3A**). Treatment with STAT3/5 inhibitors resulted in a significant reduction in tumor growth, compared to vehicle treatment. On average, MV4-11 tumor volume was reduced by 50% with JPX-0750, whereas SNK-6 tumor volume was reduced by 40% at JPX-0700 treatment end, compared to vehicle treated mice (**Figure 3B, C, E, F**). Tumor lysates were immunoblotted and probed for total STAT3 and STAT5 levels. Total STAT5 levels were reduced by 40% in AML tumors from inhibitor treated mice and by 30% in NKCL tumors, whereas total STAT3 protein levels were reduced by 60% in NKCL xenografted tumors (**Supplementary Figure S3A, B**). Endpoint analysis revealed decreased Ki-67 index and increased signals of cleaved Caspase-3 in the tumor tissue, displaying reduced cell growth and induction of cell death in JPX-0700 or JPX-0750 treated mice (**Figure 3D, G**). Importantly, we did a comprehensive toxicological analysis in drug treated mice and detected no statistically significant loss of body weight or obvious defects in overall hematopoiesis (**Supplementary Figure S3C, D, E, F**). The white blood cell count, hematocrit and platelet levels were overall unchanged. The compounds did not cause morphological changes to the liver, kidney and heart as visualized by histological staining, and liver/kidney damage parameter analysis indicated no toxic side effects (**Supplementary Figure S3G, H, I, J**). Thus, we conclude that the compounds are effective in inhibiting AML and NKCL tumor growth *in vivo* and are well tolerated in mice with no adverse toxicity.

**Figure 3.**
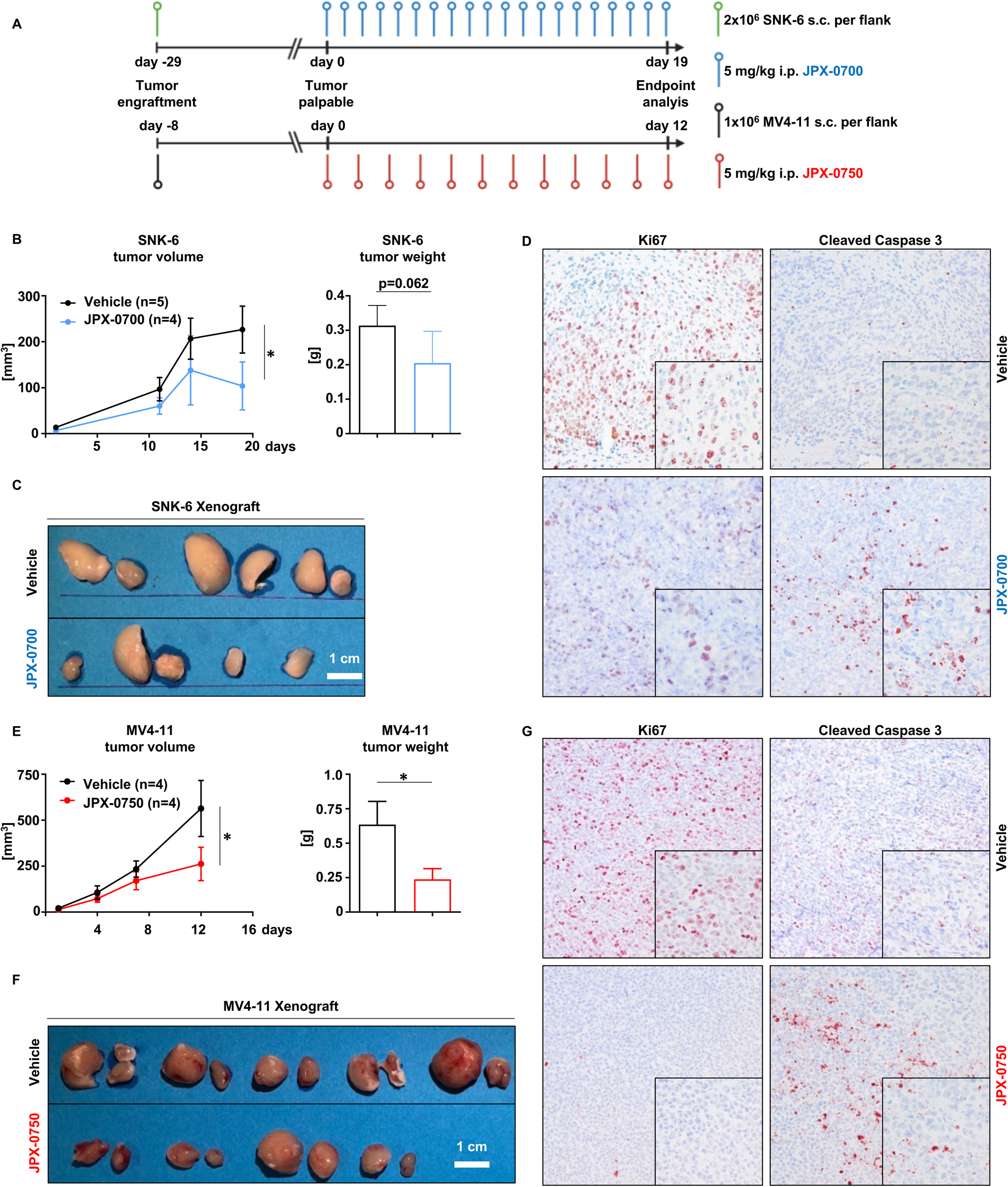
JPX-0700 and JPX-0750 can significantly block AML and NKCL tumor growth and induce growth arrest and cell death in xenografted NSGS mice. A, Summary of AML and NKCL xenograft generation. 1x106 MV4-11 or 3x106 SNK-6 cells were subcutaneously injected into NSGS mice. After the tumors became palpable at around 8 (MV4-11) or 29 (SNK-6) days post injection the mice were subjected to daily i.p. injections with 5 mg/kg JPX-0700 (19 days, NKCL xenograft) or JPX-0750 (12 days, AML xenograft). Tumor volumes and weights of B, SNK-6 and E, MV4-11 xenografted mice. Photographs of C, SNK-6 and F, MV4-11 xenografted tumors. D, SNK-6 and G, MV4-11 tumors exhibited decreased cell proliferation and viability as shown by reduced staining of Ki-67 and cleaved Caspase-3 in immunohistochemistry analysis after treatment with STAT3/5 inhibitors at the endpoint analysis of 12 (AML) or 19 days (NKCL). Data represent the mean ± SEM. *p < 0.05, by 2-way ANOVA with Bonferroni’s correction.

### The combination of dual pharmacologic STAT3/5 inhibition with standard of care antineoplastic drugs achieves synergistic effects in AML and NKCL cells

Due to the aggressiveness and heterogeneity of AML and NKCL, combinatorial therapies, if tolerable, are likely to overcome resistance and lead to long-term disease control and/or improve the quality of life of patients. To identify synergies between the STAT3/5 inhibitors and standard of care drugs for both AML and NKCL (27, 28), we tested JPX-0700 or JPX-0750 in combination with DNA-damaging drugs (doxorubicin, daunorubicin, cytarabine), cytostatic agents (methotrexate, L-asparaginase), FLT3-inhibitors (gilteritinib, midostaurin), HDAC inhibitors (vorinostat, belinostat), a multikinase inhibitor (cabozantinib), a DNA-methyltransferase inhibitor (azacitidine), and a BCL-2/xenolytic drug (venetoclax) in AML and NKCL cells *in vitro*. In general, we observed JPX-0750 having stronger synergistic effects with antineoplastic drugs in MV4-11 and MOLM-13 cells, whereas JPX-0700 demonstrated superior synergy in SNK-6 and NK-YS cells (**Figure 4A, Supplementary Figure S4A**). When combined with JPX-0750, doxorubicin, daunorubicin and cytarabine displayed the highest synergy scores, whereas FLT3 inhibitors generally showed weaker synergies with JPX-0750 in AML cells. The highest synergistic effects were observed in NKCL cells, when doxorubicin, vorinostat, or cabozantinib were combined with JPX-0700.

**Figure 4.**
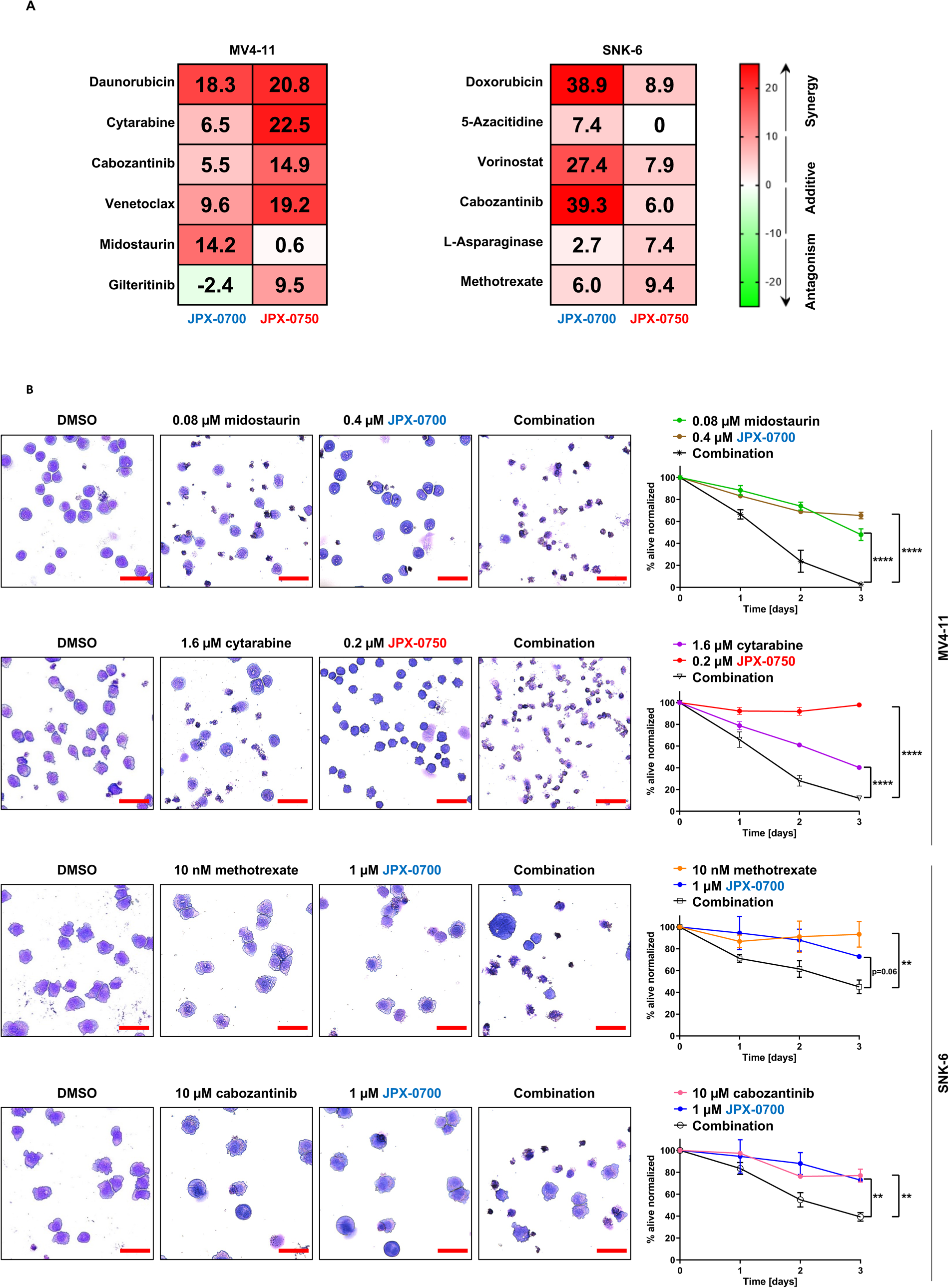
The STAT3/5 inhibitors exhibit synergy with standard of care antineoplastic drugs and targeted therapies in AML or NKCL cell lines. A, Synergy analysis of the indicated two-drug combinations in the MV4-11 and SNK-6 cells after 72 h treatment (N=2). The Zero interaction potency (ZIP) model was applied to quantify the degree of synergy. The most synergistic area (MSA) was calculated, which represents the most effective 3x3 dose-window. An MSA/ZIP score between -10 and 10 indicates that 2 drugs are additive, while a score above 10 indicates a synergistic effect. B, MV4-11 and SNK-6 cells were subjected to drug treatments, either individually or in combination, for a total duration of 72 hours. Cytospins were prepared from these cells, stained with H&E, and cell viability was assessed at 24, 48, and 72 hours following the treatment (N=2). Red bars represent 20 μm. Data represent the mean ± SEM. *p < 0.05, **p < 0.01, ***p < 0.001, and ****p < 0.0001, by 1-way ANOVA with Bonferroni’s correction.

For a better visualization of synergistic effects, cytospins were prepared from MV4-11 and SNK-6 cells treated with selected drugs in single treatments and in combination (**Figure 4B**). Two drug combinations per cell line were chosen: one drug being a first-line treatment commonly used for the disease and the second targeting a genetic abnormality present in the respective cell line. In this case, FLT3-ITD-driven MV4-11 cells were treated with cytarabine or midostaurin, while REL-mutated SNK-6 cells were treated with methotrexate or cabozantinib, in combination with the STAT3/5 inhibitors. The choice between JPX-0700 or JPX-0750 as a companion drug was based on the synergy scores in previous experiments. After 72 h of combinatorial treatment, we observed that cells were reduced in size, and cell membrane blebbing and disintegration was visible, indicating cell death, whereas single drug treatments did not lead to such significant changes in cell morphology (**Figure 4B**). In conclusion, the JPX class of compounds exhibited remarkable *in vitro* synergies when combined with clinically approved drugs, particularly JPX-0700 in NKCL and JPX-0750 in AML. We identified both inhibitors as highly effective combinatorial partners for inducing AML and NKCL cell death.

### STAT3/5 inhibitors effectively kill primary AML blasts and can be combined with approved chemotherapeutics to achieve synergistic effects

To test the translational potential of these JPX-inhibitors, we focused on the treatment of primary AML cells. We were unable to obtain viable NKCL patient samples due to their rarity and the manner in which patient tumor samples are commonly processed after resection. AML blasts from 20 patients and healthy bone marrow cells from five donors were cultured, treated for 48 h with different concentrations of STAT3/5 inhibitors, and cell viability was determined. Remarkably, the administration of both STAT3/5 inhibitors as a single agent resulted in a significant inhibition of AML blast growth, indicating their potent anti-leukemic effect (**Figure 5A**). Both JPX-0700 and JPX-0750 exhibited equivalent potency, and the mutational status of patient samples did not have a significant impact on the sensitivity observed, underlining the importance of STAT3/5 for AML survival and proliferation. Importantly, our findings demonstrate that AML blast cells exhibit a 4- and 10-fold higher sensitivity towards JPX-0700 and JPX-0750, respectively, in comparison to healthy bone marrow cells, revealing a therapeutic window (**Figure 5B**). Strikingly, when JPX-0750 was administered together with approved chemotherapeutics, including DNA damaging chemotherapies, DNA demethylating agents, BH3-mimetics, HDAC-, JAK-, MDM2- or FLT3 inhibitors, synergistic effects were observed with venetoclax, idarubicine or cytarabine (**Figure 5C**). These combinations resulted in enhanced cell killing compared to single-agent therapy.

**Figure 5.**
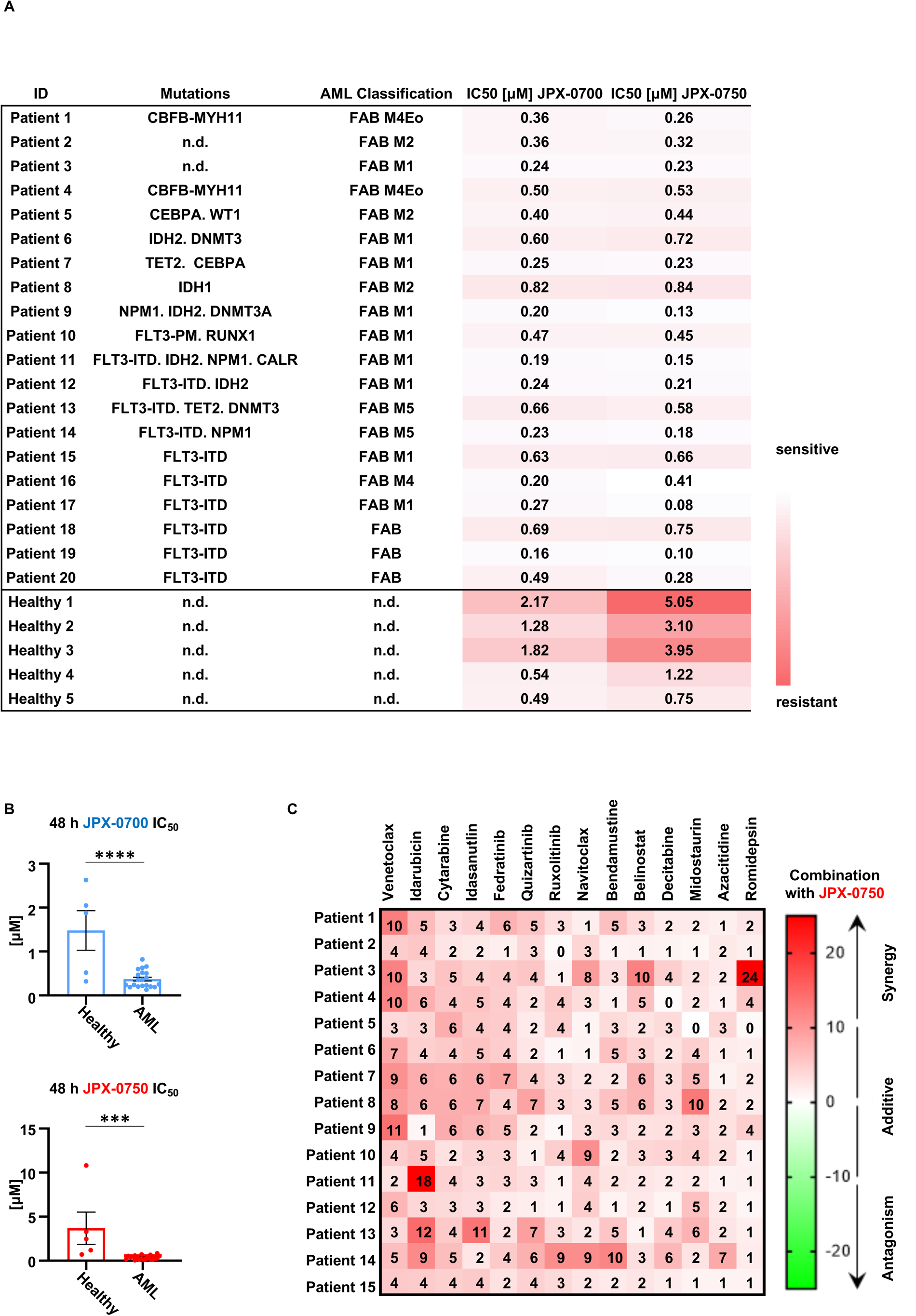
STAT3/5 inhibitors inhibit primary AML blast growth and exhibit synergies with standardly used chemotherapeutic drugs. A, AML patient characteristics, including mutations, disease staging, and 48 hour IC50 values of JPX-0700 and JPX-0750 are shown. B, Statistical analysis of IC50 values. Healthy bone marrow was used as control. C, Synergy analysis of drug combinations with JPX-0750 in AML blasts after 48 h treatment. Data represent the mean ± SEM. *p < 0.05, **p < 0.01, and ***p < 0.001, by 1-way ANOVA with Bonferroni’s correction.

Taken together, our results demonstrate the effective inhibition of primary AML blast survival and proliferation by dual STAT3/5 inhibition as mono- and combinatorial therapies. These findings highlight the potential of utilizing STAT3/5 inhibitors in combination with existing chemotherapeutic agents as a promising therapeutic strategy for AML, warranting further development of clinical grade dual-STAT3/5 inhibitors.

## Discussion

Here we investigated the efficacy of novel, dual STAT3/5 inhibitors JPX-0700/0750 in AML and NKCL. These electrophilic STAT3/5-targeting inhibitors facilitate degradation of both phospho- and total STAT3/5. Dual degradation of STAT3/5 was highly efficacious in inducing cell death and cell cycle arrest in both AML and NKCL cell lines, displaying *in vivo* anti-tumor efficacy whilst being well tolerated by xenografted mice, and demonstrating therapeutic activity against *ex vivo* AML patient samples. Moreover, the compounds displayed synergistic effects in combination with standard-of-care drugs in AML and NKCL, promoting more efficient killing of tumor cells as well as primary AML patient blasts.

Hyperactivation of STAT3/5 proteins is found in many blood cancers, with oncogenic STAT3/5 signaling promoting cancer initiation, maintenance and progression. Degradation of both STAT3/5 is an intriguing yet underexplored concept in cancer treatment, which could represent a promising therapeutic strategy for different blood cancer types. The efficacy of STAT3/5 inhibitors in both AML and NKCL might be best explained by the common mechanisms of cell cycle progression and survival driven by STAT3/5 in many blood cancers. Resistance mechanisms emerging during FLT3-ITD or FLT3-TK TKI therapy in AML include enhanced STAT3/5 expression and activation, in addition to AXL induction, enhanced D-type Cyclin or anti-apoptotic BCL-2 family member action (all targets of STAT5 and/or STAT3) (29).

STAT3/5 oncogenic signaling is likely further amplified in NKCL as a result of genetic aberrations. Through broad genetic analysis in NKCL, we found: (i) potential higher sensitivity to activating cytokines (amplified γ_c_ gene dosage, as exemplified by CN gains on *Chr X*), likely triggering JAK1/3-STAT3/5 kinase action; (ii) CN gains in *MDM2*, potentially blocking c-CBL, the E3 ligase degrading STAT5 (30); (iii) CN gains or activating GOF mutations in *STAT3* and *STAT5A/B,* boosting their transcriptional functions; (iv) CN gains in epigenetic regulators (*DNMT3A/B*, *HDAC8*); (v) gains in anti-apoptotic BCL2 family members (*MCL1*, *BCL2L1*); (vi) gains in proliferative oncoproteins (*C-*/*N-MYC*) or (vii) CN loss and LOF mutations in negative regulators (*TP53*, *CDKN2A*, *PTPN3*). *STAT3/5* CN gains together with higher γ_c_ signaling, amplified c-MYC or n-MYC, increased BCL-2/-x_L_ levels, gains in *HDAC8* (catalyzing acetylation) and gains in *DNMT3A/B* (enhancing DNA methylation) could benefit growth and survival of NKCL cells. Similarly, negative regulator loss in the context of STAT3/5 signaling revealed mutations in *PTPN3*, and *TP53/CDKN2A* tumor suppressors, in the majority of NKCL cell lines. Overall, this suggests that a general cytokine receptor-JAK1/3-STAT3/5 driver axis could exist in AML and NKCL, synergizing with *MYC* family member amplification, loss of p53/p16^INK4A^, as well as enhanced DNA methylation. Lost p53/p16^INK4A^ is consistent with overcoming senescence and failure to arrest upon accumulation of reactive oxygen species (ROS)-mediated DNA damage. Enhanced ROS levels also inhibit phosphatase action, and several key phosphatases displayed CN loss in NKCL. *TP53* is frequently deleted or mutated in leukemia/lymphoma (9 out of the 14 cell lines analyzed here), but the high frequency of *TP53* genetic alterations seen in the cell lines is not observed in patient samples, which could represent clonal outgrowth in culture conditions (9). Importantly, it is the mutational landscape and, as a consequence, changed interactions with oncogenes or tumor suppressors, together with epigenetic chromatin remodeling, that influence whether STAT3/5 act as oncogenes or promote senescence or trans-differentiation (31).

Degradation-based therapeutic strategies offer superior potential. We profiled two drugs from this class of compounds that displayed efficacy in degrading STAT3/5 proteins in both AML and NKCL. These degraders can also block non-canonical protein functions, thus eliminating non-enzymatic/activity-based scaffolding or cofactor/corepressor recruitment as well as mitochondrial STAT3/5 function (32), which was shown to be essential for oncogenic transformation mediated by RAS/RAF (12). Non-phosphorylated STAT5A reportedly stabilizes heterochromatin formation by interacting with Heterochromatin Protein alpha, balancing oncogenic phosphorylated STAT5A/B activity (33). Studies have also revealed that STAT5A/B interact with chromatin remodelers to influence chromatin organization at promoter and enhancer regions via oligomerisation/DNA looping, similarly to Cohesin ring formation (34). Dual STAT3/5 degraders highlight the potential importance of blocking broader STAT functions beyond those mediated by tyrosine phosphorylation, affecting a larger spectrum of important cancer cell processes. STAT proteins possess a long half-life, but they paradoxically express a low intrinsic thermodynamic stability, as shown by low melting temperatures (35). Interestingly, STAT3/5 engagement with JPX compounds *in vitro* has been reported to further reduce the melting temperature and is hypothesized to lead to a non-functional protein, disrupting protein-protein interactions, and rendering STAT3/5 susceptible to the cellular degradation machinery (23). The primary concern for AML or NKCL treatment lies in achieving effective drug penetration into the bone marrow or achieving sufficient penetration into the skin, lymph nodes, blood vessels, and mucosal tissues. As such, JPX-0700/0750 demonstrated a strong reduction in leukemic burden *in vivo* at low doses of 5 mg/kg (IP, once daily) which is only 2-5% of the dosage employed with PROTACs highlighting therapeutic value of monovalent degraders (36). Thus, lower molecular weight compounds would be predicted to be more efficacious as a result of more effective penetration through tissue barriers and lower dosage required to inhibit leukemic burden, as shown in our successful xenograft experiments here.

AML patients >60 years of age on standard treatment have a median overall survival time of <1 year, despite that several novel targeted drugs have received clinical approval in the recent past. Even worse, stage IV NKCL patients have a median survival time of <4 months (37–39). Many AML/NKCL patients do not achieve long-term remission, and chemotherapy often has severe side effects compromising quality of life. Due to the high prevalence of mutations in epigenetic modifiers, chromatin remodeler proteins, signaling TK and cell cycle regulators in both blood cancer types, the use of a single agent is insufficient to eradicate cancer (19, 27). Our drug synergy results using clinically approved therapeutics together with JPX-0700/0750 highlight the possibility to overcome drug resistance. While doxorubicin, a non-specific chemotherapeutic and cabozantinib, a multikinase inhibitor, highly synergized with JPX inhibitors in both AML/NKCL, the epigenetic drug azacitidine and the BCL-2 inhibitor venetoclax had differential impact on these diseases. Different synergy outcomes could be explained by the presence of CN gains in the *DNMT3A*/B and *BCL2L1* locus in SNK-6 cells, decreasing efficacy of azacitidine and venetoclax (40). We also noted a limited level of synergy when combining FLT3 TKI (midostaurin and gilteritinib) with the STAT3/5 compounds in FLT3-ITD^+^ AML cell lines. These findings align with the notion that drug combinations are more effective when the targeted signaling pathways are more independent/distinct (41). The synergistic effect of drug combinations with STAT3/5 inhibitors, specifically in FLT3-ITD-driven AML, could have merit in patients facing resistance to TKI therapy (40, 42). Venetoclax in combination with epigenetic enzyme blockers leads to a metabolic rewiring in leukemic stem cells, and studies have revealed that STAT3/5 can act in non-canonical ways to influence chromatin organization as well as metabolism (12, 40).

In summary, the effectiveness of dual inhibition of phospho-and total STAT3/5 by JPX inhibitors in AML/NKCL emphasizes their essential roles in initiating and driving these cancers. Our findings provide a rationale for the clinical development of these compounds towards their combinatorial use for blood cancer treatment.

## Materials and Methods

### Cell culture and generating stable cell lines

Ba/F3, MOLM-13, MV4-11, KHYG-1, SU-DHL-1, NK92, YT, NHDF, A549 and Mac-2A cell were obtained from the German Collection of Microorganisms and Cell Cultures (DSMZ, Germany). NKL and MTA cells were kindly provided by Dr. Raphael Koch (University Medical Center Göttingen, Germany). SNK-6 and NK-YS cells were provided by Dr. Wing C. Chan (City of Hope Medical Center, Duarte, USA). My-La and HUT78 cells were provided by Dr. Marco Herling (University of Leipzig, Germany). Mac-2A cells were provided by Marshall Kadin (Boston University, USA). The authenticity of the cell lines was confirmed by analysis of highly polymorphic short tandem repeat loci (STR) using the PowerPlex 16 HS System (Promega; performed by Microsynth AG, Switzerland). All cell lines were maintained in RPMI-1640 (Gibco) supplemented with 10% FBS (Biowest), 0.06 g/L penicillin/0.1 g/L streptomycin (Pen/Strep, VWR), and 2 mM L-glutamine (Life Technologies). Culture media of NK92, NKL, SNK-6, NK-YS, KHYG-1 cells was additionally supplemented with 5 ng/mL recombinant human IL-2 (ImmunoTools GmbH, Germany). Ba/F3 cells were grown in the presence of 1 ng/mL murine IL-3 (mIL-3, Immunotools). Mycoplasma contamination was regularly excluded using the MycoAlert mycoplasma detection kit (Lonza Group AG, Switzerland). All cell lines were cultured at 37°C in a humidified atmosphere containing 5% CO_2_. Experiments were performed within 20 passages after cell resuscitation. None of the above-mentioned cell lines are listed in the register of cell lines that are known to be misidentified through cross-contamination. Mammalian expression constructs containing FLAG-tagged human STAT3, STAT5A or STAT5B in a pMSCV-IRES-GFP plasmid were used to generate stable Ba/F3 cell lines. The procedure of retroviral transduction with these constructs was performed as previously described (43).

### Immunoblot analysis

Approximately 0.75x10^6^ cells/mL were seeded into a 6-well plate (Corning) and treated with the desired concentration of the drug of interest, or the appropriate cytokine, the following day. Cells were incubated at 37°C for 24 h and then lysed in immunoprecipitation assay (IP) buffer [25 mM HEPES (pH 7.5), 25 mM Tris-HCl (pH 7.5), 150 mM NaCl, 10 mM EDTA, 0.1% Tween-20, 0.5% NP-40, 10 mM β-glycerophosphate] supplemented with cOmplete, EDTA free Protease

Inhibitor Cocktail (Roche), 1 mM sodium orthovanadate, 1 mM NaF, 10 μg/mL Leupeptine, 10 μg/mL Aprotinine and 1 mM PMSF. Protein concentrations were determined by the Bradford protein assay. Aliquots of total cell lysates were fractionated on SDS polyacrylamide gels and transferred to nitrocellulose membranes (Cytiva). Intercept (TBS) Blocking Buffer (LI-COR Biosciences) was used for blocking and preparing the antibody dilutions. Information about the antibodies and their dilutions used for Western blotting is available in Table 1. Equal loading was confirmed by probing the same membranes with a specific antibody for human HSC70, ACTIN or α-TUBULIN.

**Table 1.** Antibodies used for immunoblotting experiments.

| Target | Catalog # | Supplier | Species | Dilution |
| --- | --- | --- | --- | --- |
| ACTIN | 1615 | Santa Cruz Biotechnology | rabbit | 1:5000 |
| CDK6 | 177 | Santa Cruz Biotechnology | rabbit | 1:1000 |
| CC3 | 9661 | Cell Signaling Technology | rabbit | 1:1000 |
| c-MYC | ab32072 | Abcam | rabbit | 1:1000 |
| CYCLIN D2 | 3741S | Cell Signaling Technology | rabbit | 1:1000 |
| CYCLIN D3 | 2936T | Cell Signaling Technology | mouse | 1:1000 |
| HSC70 | 7298 | Santa Cruz Biotechnology | mouse | 1:5000 |
| PARP | 9532S | Cell Signaling Technology | rabbit | 1:1000 |
| PIM1 | 374116 | Cell Signaling Technology | rabbit | 1:1000 |
| p-Tyr-STAT3 | 9131S | Cell Signaling Technology | rabbit | 1:1000 |
| p-Tyr-STAT5 | 9314S | Cell Signaling Technology | rabbit | 1:1000 |
| STAT3 | 610189 | BD Biosciences | mouse | 1:1000 |
| STAT5 | 610191 | BD Biosciences | mouse | 1:1000 |
| α-TUBULIN | 32293 | Santa Cruz Biotechnology | mouse | 1:5000 |

### Cell viability assays

For each assay, 10000 cells/well were seeded in triplicates on a flat-bottom 96-well sterile culture plate with low-evaporation lids (Costar). The following day, cells were treated with serial 2-fold dilutions of the drug of interest, or 0.5% DMSO used as vehicle control, which was also the starting/maximum DMSO concentration when diluting the drugs. The compounds were manually transferred to plates containing cells. Bortezomib (MedChemExpress) [50 µM] was used as a positive control. Treated cells were incubated at 37°C and 5% CO_2_ for 72 h. Cell viability was measured using the CellTiter-Blue Cell Viability Assay (Promega) and the GloMax® Discover Microplate Reader (Promega). IC_50_ values were calculated using non-linear regression of log-transformed, normalized data in GraphPad Prism 7.0 (GraphPad Software, Inc.). Three independent experiments were performed.

### Oligonucleotide array comparative genomic hybridization (aCGH)

DNA extraction on cell lines was performed using the QiaAmp DNA Blood Mini kit (Qiagen). Reference DNA for the analysis was derived from multiple healthy and anonymous male donors (Promega). The labeling and hybridization steps were conducted in accordance with the provided instructions using the SureTag DNA Labelling kit (Agilent Technologies, USA). Equal amounts of DNA labeled with fluorescent markers were combined and co-hybridized onto a 44LK DNA microarray. (G4426B-014950, Agilent Technologies). Following hybridization and washing, in adherence to the manufacturer’s guidelines, scanning was conducted using the G2505B Micro Array Scanner (Agilent Technologies). Subsequently, Feature Extraction (version 10.7.3.1) and Agilent Genomic Workbench version 7 were employed for data analysis. The analysis utilized the ADM-1 algorithm with a threshold of 6, and aberration boundaries were defined as ±0.25 (log2 ratio) with a minimum of three probes per region, as recommended by Agilent Technologies.

### Electrophoretic mobility shift assay

Single stranded DNA oligos (see Table 2) were purchased from Integrated DNA Technologies and were annealed under normal PCR conditions. Oligo labeling and electrophoretic mobility shift assay (EMSA) was performed for STAT5A/B using the β-casein response element (1xβCas) and for SP1 using a GC-rich motif (bSP1) (44). For STAT5A supershifts, a polyclonal rabbit sera against the C-terminus (NH2-LSPPAGLFTSARSSLS-COOH, Eurogentech) and for STAT5B supershifts, the G-2 antibody (sc-1656, Santa Cruz Biotechnology) was used. Detection of gels was facilitated by a phosphorimager-detection system (GE Healthcare).

**Table 2.**
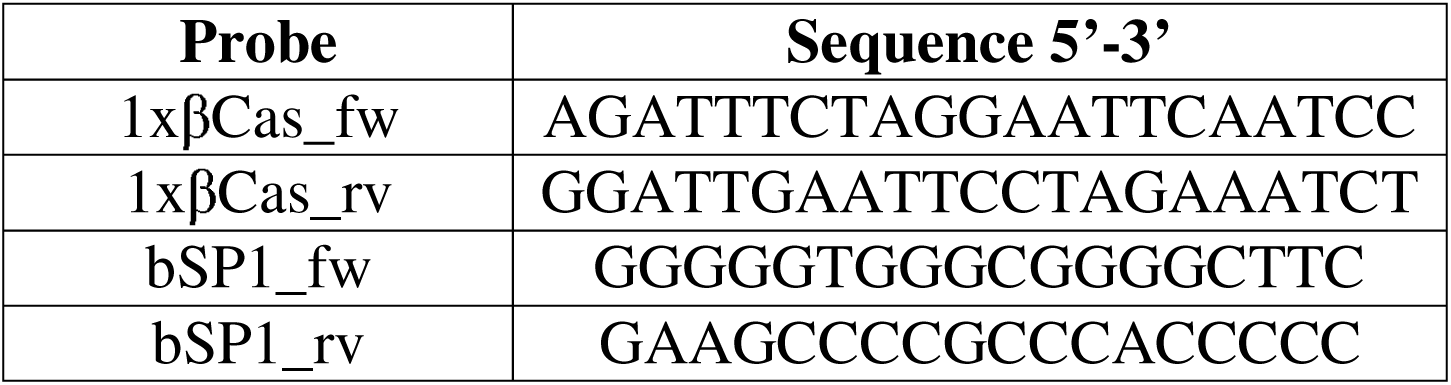
DNA probes used for EMSA experiments.

### Cell death and cell cycle measurements

The effect of the drugs used was assessed at 24 h post treatment using flow cytometric analysis of Annexin V and propidium iodide staining according to the manufactureŕs instructions (eBioscience). Samples were analyzed on a FACSCanto II analyzer and data analysis was completed using FlowJo 7.6 software (BD Biosciences). Bortezomib (5 µM) was used as a positive control. The procedure of cell cycle staining was performed as previously described (10).

### Patient Samples

Bone marrow cells were extracted from the iliac crest of patients with AML and were collected and stored at -80°C until used. The study was approved by the ethics committee of the Medical University of Vienna (1184/2014 and 1334/2021) and the Helsinki University Hospital (303/13/03/01/2011) and was conducted in accordance with the declaration of Helsinki. All patients gave written informed consent. Diagnoses were established according to French-American-British (FAB) and World Health Organization (WHO) criteria. Cells were maintained in RPMI 1640 supplemented with 10% FBS, 10 U/mL penicillin, 10 μg/mL streptomycin, 2 mM L-glutamine. Viability of the samples was normalized to the vehicle (DMSO) control.

### In vitro synergy screen

The analysis was performed using the SynergyFinder 3.0 web application (45). The impact of the drug combinations on leukemia and lymphoma cell lines was calculated using synergy scores within the most synergistic concentration window (3x3), which allow for the comparison of drugs at their most beneficial concentration ranges (45). The degree of synergy was quantified using the Zero interaction potency (ZIP) model. The ZIP model assumes that two non-interacting drugs are expected to incur minimal changes in their drug response curve, combining the advantages of Loewe additive model and Bliss independence model. The four-parameter logistic regression (LL4) was used as the curve-fitting algorithm. The cNMF algorithm was applied to detect and replace outlier measurements when necessary (46). A ZIP synergy score less than -10 indicates when the interaction between two drugs is likely to be antagonistic; from -10 to 10 indicates when the interaction between two drugs is likely to be additive; a value larger than 10 indicates that the interaction between two drugs is likely to be synergistic. All assays were repeated in at least two biological replicates.

### Ex vivo synergy screen

Primary cells were cultured in Mononuclear Cell Medium (#C-28030, PromoCell) supplemented with 2.5 ug/ml amphotericin B (#A2942, Sigma-Aldrich) and 10 ug/ml gentamicin (#15710-049, Thermo Fisher). Drug sensitivity testing was carried out at the FIMM High Throughput Biomedicine Unit, which is hosted by the University of Helsinki and supported by HiLIFE and Biocenter Finland. Drug sensitivity testing was performed as previously described (47–49). Briefly, drugs were dispensed on 384-well plates (#3712, Corning) using an Echo Liquide Handler (Labcyte). To dissolve the drugs, 5 μL/well of Mononuclear Cell Medium (#C-28030, PromoCell) were dispensed using a MultiFlo FX dispenser (BioTek) and after that 10 000 cells in 20 µL of Mononuclear Cell Medium were dispensed on the plates. After incubating the cells for 48 hours at 37 °C and 5 % CO2, the cell viability was assessed using the CellTiter-Glo (CTG) luminescent assay (Promega) and a Pherastar FS plate reader (BMG Labtech). To calculate the relative efficacy (% inhibition), 100 μM benzethonium chloride was used as a positive (total killing) and 0.1 % DMSO as a negative (non-effective) control. To quantify the efficacy each compound, a drug sensitivity score (DSS) was calculated as previously described (50). In pairwise drug combination testing, 1:1 combination of seven doses for each drug was used. After that, the DECREASE model predictions were used to fill the full (8L×L8) drug combination dose-response matrices (51). The observed responses were compared with expected combinatorial responses, calculated based on the ZIP reference model, to classify drug combinations as synergistic, additive, or antagonistic (52). The ZIP model-based summary synergy scores, were calculated using SynergyFinder (45).

### In vivo efficacy and safety evaluation

The animal research was conducted following the institutional regulations for animal welfare and with the approval of the Austrian Federal Ministry of Science, Research, and Economy (BMBWF-68.205/0130). Xenografted NSGS mice were treated with JPX-0700 (5 mg/kg), JPX-0750 (5 mg/kg) or vehicle (5% ethanol, 5% Kolliphor-EL (Carl Roth), 30% propylenglycol (Carl Roth), 20% HP-ß-cyclodextrin (Sigma Aldrich) in PBS) by intraperitoneal injection daily for 12 or 19 days. Blood was obtained by heart puncture and collected in EDTA-tubes (Mini-Collect K_3_EDTA tubes). Blood parameters were measured using an animal blood counter (Vet abc, scil animal care). Plasma was prepared by centrifugation of the whole blood for 20 min at 5000 rcm. Serum concentration of aspartate aminotransferase (AST), alanine aminotransferase (ALT) and blood urea nitrogen (BUN) was measured using a chemistry analyzer (IDEXX VetTest 8008,

IDEXX Laboratories). Tumor volumes were calculated using the formula: (tumor length x tumor width^2^): 2. Organs were fixed overnight in 4% phosphate buffered formaldehyde solution (Roti® Histofix, Carl Roth), dehydrated, paraffin embedded and cut into 2 µm consecutive mouse organ or tumor sections. Sections were stained with Hematoxylin (Merck), Eosin G (Carl Roth), Ki67 (MM1-L-CE, Leica Biosystems) and Cleaved Caspase 3 (#9664, Cell Signaling Technology).

### Statistical analysis

GraphPad Prism 7 and 8 were used to perform the statistical analyses. Statistical tests are specified in Figure legends. The threshold for statistical significance was set to P < 0.05, unless otherwise specified. P-value: <0.05 (*), <0.01 (**), <0.001 (***), <0.0001 (****).

## Data and materials availability

All data associated with this study are present in the paper or in the supplementary materials.

## Acknowledgments

We wish to thank Safia Zahma and Michael Machtinger from the Institute of Animal Breeding and Genetics at the University of Veterinary Medicine Vienna for technical support in histology and during in *vivo* experiments.

## Author contributions

Conceptualization: DP, HS and RM

Resources: RM

Data curation: DP and HS

Software: HS, DP, AO, AS, MS, EDA, SHT, AI, HK, CP

Formal Analysis: HS, DP, AO, AS, HAN, EDA

Supervision: RM

Funding acquisition: RM

Validation: DP and HS

Investigation: DP and HS

Visualization: DP and HS

Methodology: HS, DP, AS, EDA, HAN, AO, SHT, DIA, AI, HK, MS, CW, TS, MLM, MB, MD, FG, RF, CP, WB, EH, WRS, LK, PV, TA, MH, SM, PTG

Writing – original draft: DP, HS and RM

Project administration: RM

Writing – review & editing: all authors

## Supplementary Tables

**Supplementary Table S1.** Mutational data of cell lines.

| Cell line | Reference |
| --- | --- |
| MV4-11 | (53, 54) |
| MOLM-13 | (53, 55) |
| NKL | (15) |
| NK92 | (15) |
| SNK-6 | (15) |
| NK-YS | (15) |
| MTA | (15) |
| KHYG-1 | (15) |
| YT | (15) |
| SU-DHL-1 | (14) |
| Mac2A | (14) |
| My-La | (2) |
| HUT78 | (2) |

## Supplementary Figure legends

**Figure S1.**
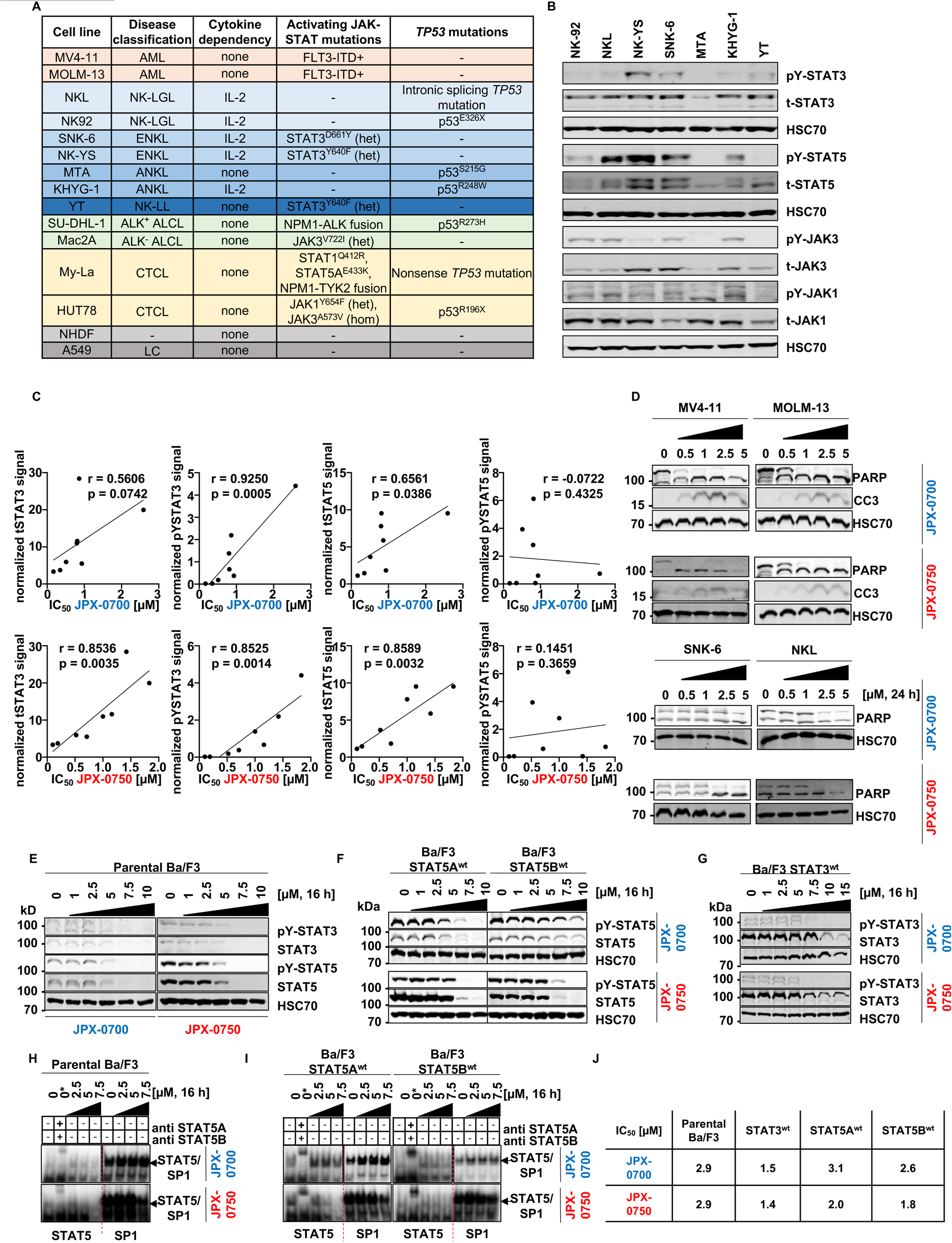
AML and NKCL cell lines harbor mutations in the STAT3/5 signaling pathway and are sensitive towards STAT3/5 degradation by JPX-0700/0750. **A,** Cell line characteristics of AML, NKCL, TCL and control cell lines **B,** NKCL cell lines were immunoblotted for total and phospho-Tyr (1022/1023)-JAK1, total and phospho-Tyr (980/981)-JAK3, total and phospho-Tyr (705)-STAT3 or total and phospho-Tyr (694/699)-STAT5A/B. HSC70 served as loading control (n=2). **C,** Pearson correlation between normalized pY/total-STAT3/5 levels and the IC_50_ values of JPX-0700 and JPX-0750 **D,** MV4-11 and SNK-6 cells were immunoblotted for PARP and Cleaved Caspase 3 (CC3) levels after 24 h treatment with JPX-0750 (N=2). **E,** Parental **F,** STAT5A^wt^ and STAT5B^wt^ or **G,** STAT3^wt^ overexpressing Ba/F3 cell lines were treated for 16 h with JPX-0700 and JPX-0750, stimulated with 10 ng/mL mIL-3 30 min before harvesting and were immunoblotted for total and phospho-Tyr (705)-STAT3 and total and phospho-Tyr (694/699)-STAT5A/B. HSC70 served as loading control (N=2, representative experiments are shown). **H and I**, Isolated protein from parental, STAT5A^wt^ and STAT5B^wt^ overexpressing Ba/F3 cell lines were probed for STAT5 dimer activity with a STAT5-specific DNA-binding element. Specificity of the observed complexes was verified by supershift analysis (*). The DNA binding activity of the transcription factor SP1 on a GC-rich DNA consensus sequence was used as loading control. **J,** 72 h IC_50_ values of Ba/F3 cell lines treated with JPX-0700/0750 (N=2, representative experiments are show). Abbreviations: *ALCL=anaplastic large cell lymphoma, AML=acute myeloid leukemia, ANKL=aggressive NK cell leukemia, CTCL=cutaneous T cell lymphoma, ENKL=extra-nodal NK/T cell lymphoma, nasal type, LC=lung carcinoma, NHDF=normal human dermal fibroblasts, NKCL=NK cell lymphoma/leukemia, NK-LGL=NK large granular lymphocytic leukemia, NK-LL=NK large lymphocytic leukemia, TCL=T cell lymphoma/leukemia*

**Figure S2.**
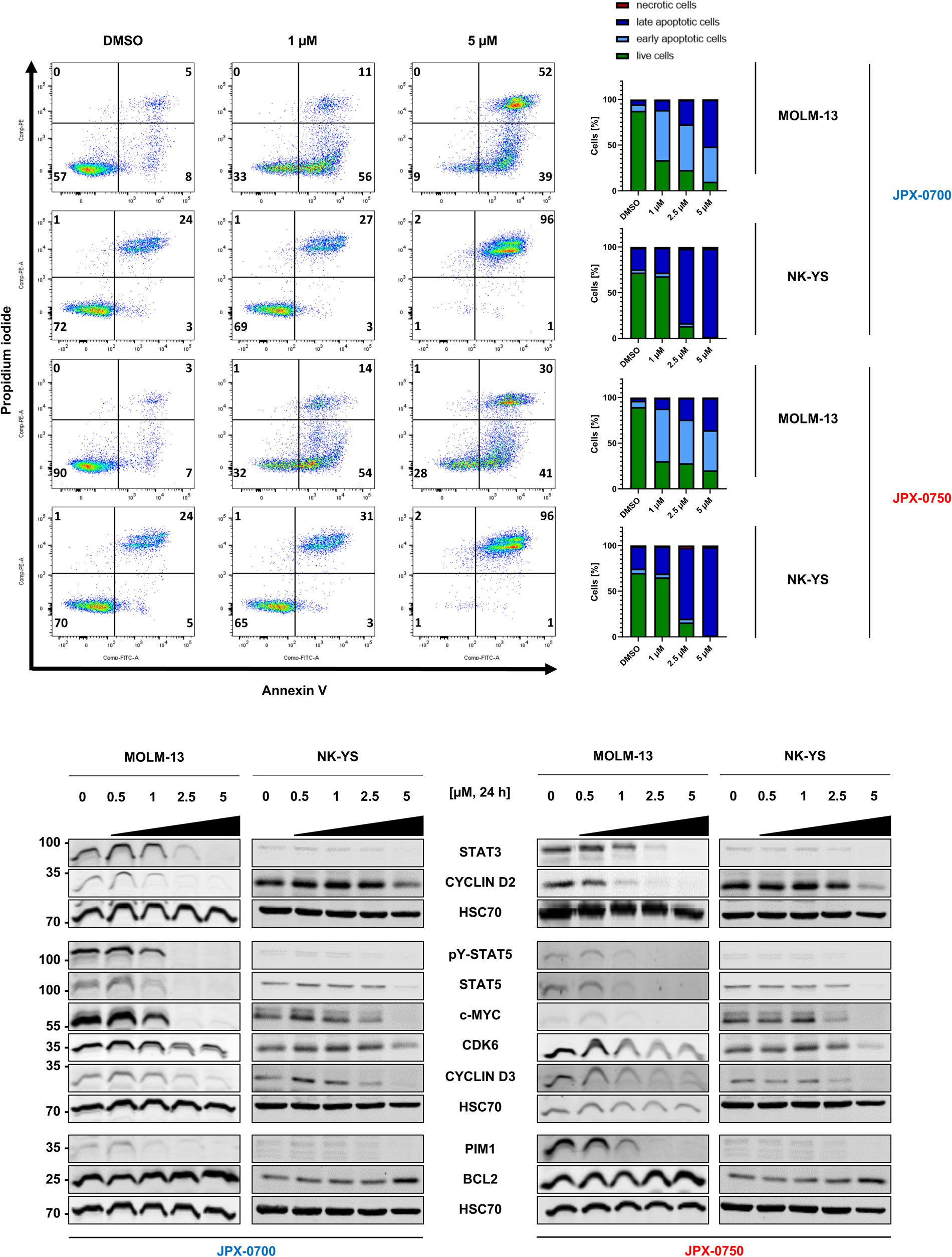
Small molecule STAT3/5 inhibitors inhibit downstream STAT3/5 signaling thereby inducing apoptosis of AML and NKCL cell lines. **A,** MOLM-13 and NK-YS cells were treated with different concentrations of JPX-0700, JPX-0750 or DMSO in a dose-dependent manner for 24 h. Apoptotic cells were detected by Annexin-V/Propidium iodide staining. Representative dot plots and quantification bar graphs are shown (N=2). **B,** Inhibitor treatment was carried out with increased dose escalation for 24 h in MOLM and NK-YS cell lines. Cells were immunoblotted for total and phospho-Tyr (705)-STAT3 and total and phospho-Tyr (694/699)-STAT5A/B. STAT3/5 target gene products c-MYC, PIM1, as well as proteins regulating the cell cycle, such as CYCLIN D2 and D3, and CDK6, were probed by Western blotting. HSC70 served as loading control (N=2).

**Figure S3.**
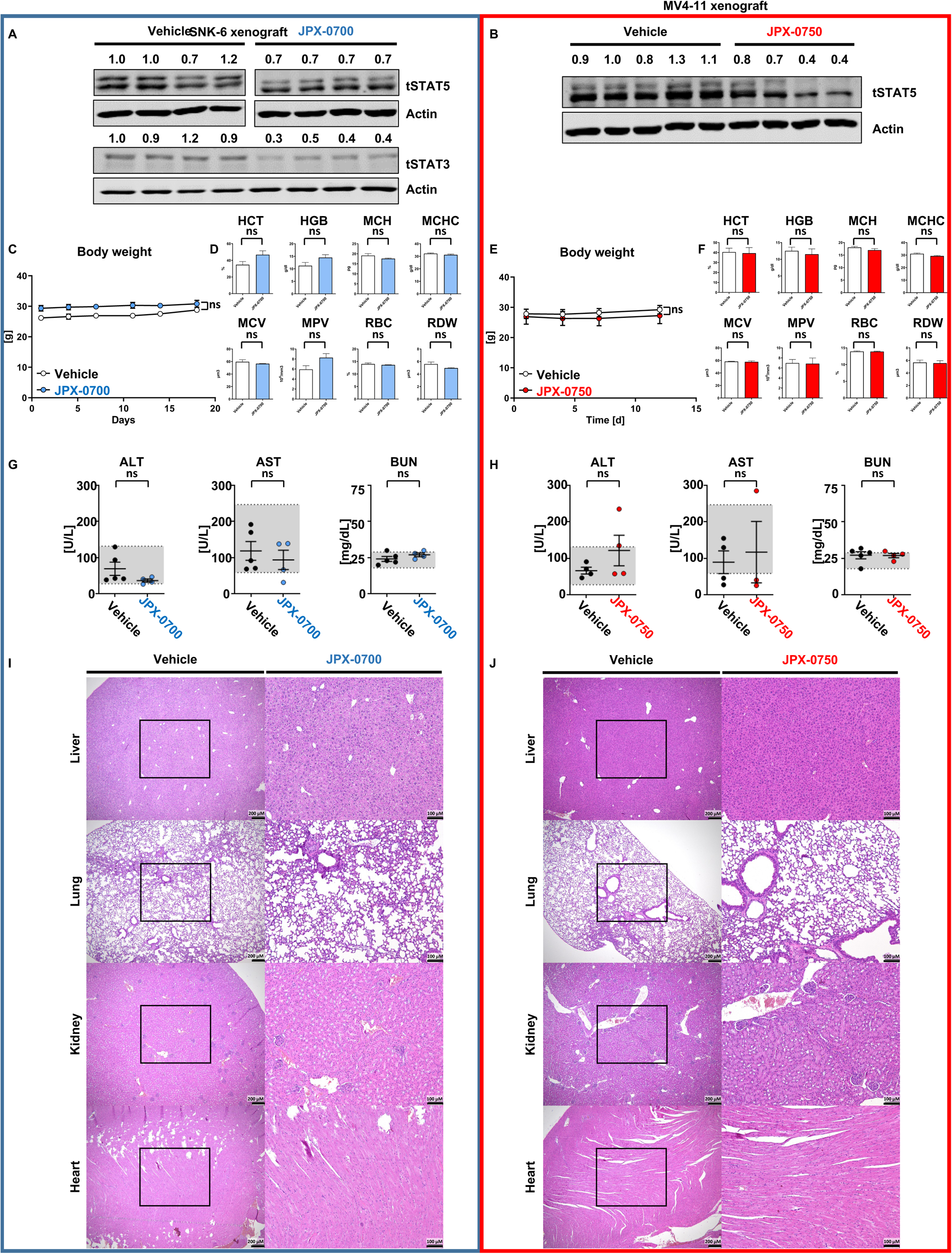
The small molecule inhibitors are efficacious in AML/NKCL xenografts and display safety *in vivo*. **A and B,** Tumors were lysed and immunoblotted for total and phospho-Tyr (694/699)-STAT5A/B and phosphor-Tyr STAT3. Actin served as loading control. **C and E,** Bodyweight during the experiment and **D and F,** Hematological parameters of xenografted mice at the endpoint. **G and H,** JPX-0700 and -0750 treated mice revealed no significant changes in Alanine Aminotransferase (ALT), Aspartate Transaminase (AST) and blood nitrogen urea (BUN), indicating no significant liver and kidney toxicity and the experimental endpoint. **I and J,** Gross organ morphology of liver, kidney, lung and heart of vehicle-, JPX-0700-or JPX-0750-treated mice at experimental endpoint. Data represent the mean ± SEM, by 1-way ANOVA with Bonferroni’s correction. *Abbreviations*: *HCT=hematocrit, HGB=hemoglobin, MCH=mean cell hemoglobin, MCHC=mean cell hemoglobin concentration, MCV=mean cell volume, RBC=red blood count, RDW=red cell distribution width, MPV=mean platelet volume*

**Figure S4.**
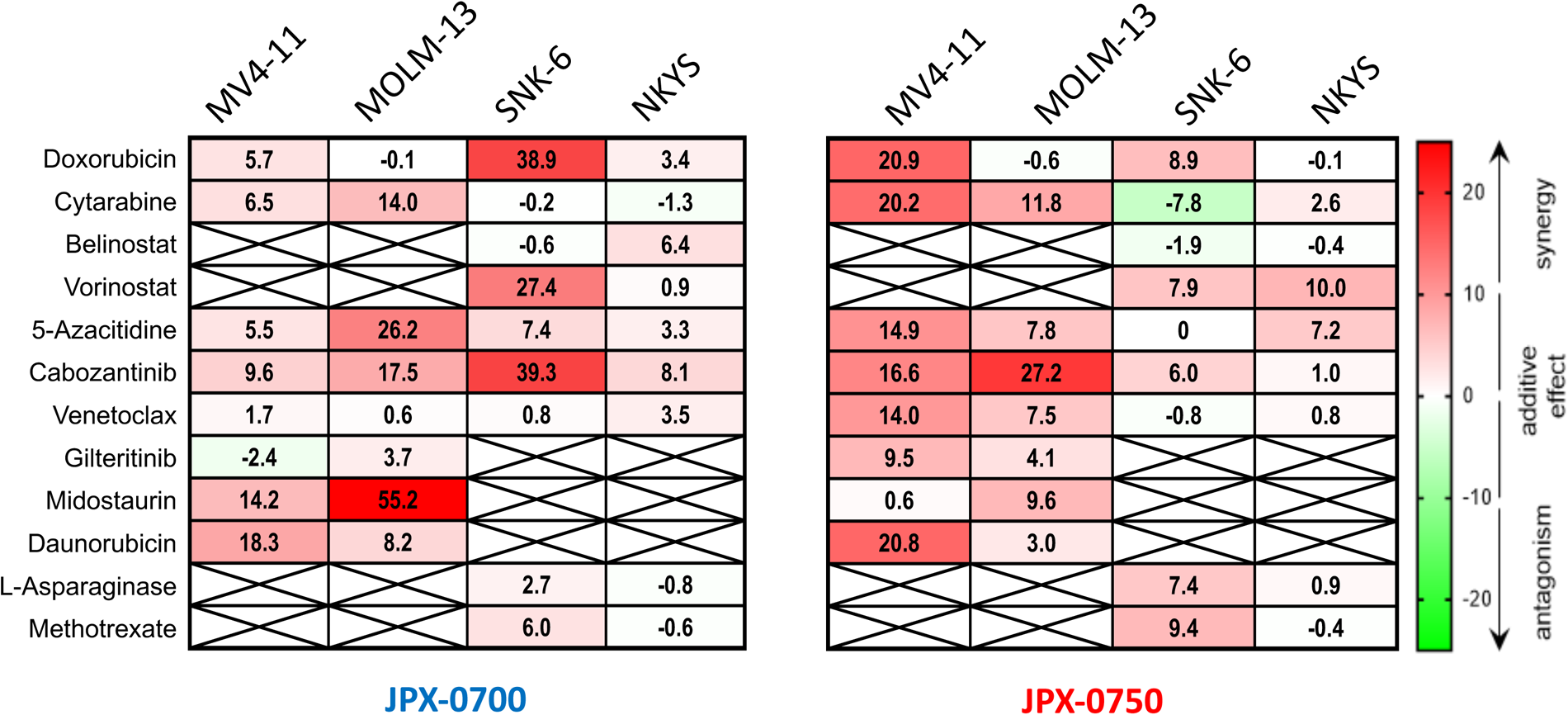
Targeted therapies, display diverse effects in combination with JPX-0700 and JPX-0750 in AML and NKCL. Synergy analysis of the indicated two-drug combinations in MV4-11, MOLM-13, SNK-6 and NK-YS cells after 72 h treatment (N=2). The Zero interaction potency (ZIP) model was applied to quantify the degree of synergy. In each graph the most synergistic area (MSA) is highlighted, which represents the most synergistic 3-by-3 dose-window. An MSA/ZIP score between -10 and 10 indicates that 2 drugs are additive, while a score above 10 indicates a synergistic effect.

